# Cortical Activity During Sustained Isometric Ankle Contractions Following Chronic Sleep Restriction: A High-Density EEG Study

**DOI:** 10.64898/2026.07.07.737078

**Authors:** Maura Seynaeve, Jessica Samogin, Dante Mantini, Toon de Beukelaar

## Abstract

**Background:** Chronic sleep restriction (CSR) impairs cognitive function, but its effects on the cortical dynamics underlying active motor performance remain poorly understood. High-density EEG provides a means to examine task-related oscillatory activity across sensorimotor and attentional networks during movement.

**Methods:** Fifteen healthy males completed a randomized crossover study involving a CSR condition (five hours sleep per night for four nights) and a control condition (normal sleep). Before and after each intervention, participants performed sustained isometric ankle contractions at 40% of their maximal force while EEG was recorded. Source-reconstructed event-related desynchronization (ERD) was computed across theta, alpha, beta, and gamma bands in the sensorimotor network and dorsal attention network. Sustained attention was assessed with the Psychomotor Vigilance Task (PVT) and perceived workload with the NASA Task Load Index.

**Results:** CSR successfully reduced sleep duration by 2.36 hours on average (p < .001). Following CSR, PVT reaction times increased significantly (Δ = +31 ms, p = .002) and attentional lapses increased (Δ = +9.87, p < .001). CSR produced a significant overall increase in ERD across bands, networks, and movement directions (F(1, 5713) = 14.20, p < .001). This effect was present in both the sensorimotor and dorsal attention networks. The ERD increase was specific to dorsiflexion and absent during plantarflexion (condition x session x movement direction: F(1, 5713) = 9.13, p = .003). Subjective mental demand increased following CSR (p = .027), while objective motor performance was largely unimpaired.

**Conclusion:** CSR increased broadband ERD during dorsiflexion across both sensorimotor and attentional networks, alongside impaired sustained attention and greater perceived mental demand. As motor performance was largely preserved, this increased ERD may reflect compensatory neural recruitment under sleep pressure.

## Introduction

Sleep is essential for maintaining optimal cognitive and physical functioning. Yet, insufficient sleep is highly prevalent in modern society, with many individuals failing to achieve the recommended 7–9 hours of sleep per night (1, 2). A common consequence is chronic sleep restriction (CSR), which occurs when small amounts of sleep are lost across multiple consecutive nights. Unlike total sleep deprivation, which is relatively rare outside of controlled laboratory settings, CSR reflects the reality of many people’s daily lives, whether due to work demands, social obligations, or lifestyle habits. CSR produces cumulative impairments in vigilance, sustained attention, executive functioning, and cognitive control that are often underestimated by those experiencing them (3, 4). These deficits are particularly evident in tasks requiring prolonged concentration, continuous response monitoring, and flexible allocation of attentional resources.

Although these functions are primarily cognitive in nature, they also contribute meaningfully to the execution of motor tasks. Motor performance is not solely determined by peripheral muscular function or primary motor execution. Sustained or precise motor tasks, such as gait control and force regulation, rely meaningfully on these higher-order cognitive processes. This is evidenced clearly in the dual-task literature, where the concurrent performance of a cognitive task reliably degrades motor output, and conversely, increased motor task demand draws on attentional and executive resources (5, 6). For instance, gait stability has been shown to deteriorate when attentional resources are divided, an observation consistent with the view that even habitual walking requires attentional control and executive function to some degree. A practical illustration of this is the well-documented reduction in movement speed when individuals engage with a mobile device while walking, reflecting an implicit cognitive recalibration to reduce fall risk (7). Cognition and motor control should therefore not be treated as independent domains; their interaction is a fundamental feature of movement at every level of complexity.

Consequently, motor tasks that rely on cognitive processes such as sustained attention and executive control may be particularly vulnerable to sleep loss. For example, sleep restriction has been shown to impair visually guided force production, with participants producing less force at the target level after four nights of restricted sleep. This is consistent with the notion that sustained attention is highly sensitive to insufficient sleep (8). However, evidence from gait control suggests that the effects of sleep loss may not be explained solely by reduced cognitive resources. In a recent study directly comparing the effects of sleep deprivation and dual-tasking on gait, sleep deprivation resulted in longer steps, reduced instantaneous lateral stability, and increased reliance on center-of-mass state during foot placement. Because both sleep deprivation and dual-tasking are thought to tax cognitive resources, overlapping gait adaptations would be expected. Yet, the gait changes observed under sleep deprivation only partially overlapped with those induced by dual-tasking, suggesting that sleep loss may impair motor control through mechanisms extending beyond a simple reduction in cognitive capacity.

Alternatively, sleep loss may impair motor performance by disrupting sensorimotor processing, including sensorimotor integration as well as feedback and feedforward control mechanisms. Postural control, a motor task that relies heavily on sensorimotor integration, has consistently been shown to deteriorate following sleep deprivation (9, 10). Notably, these sleep-related impairments in postural control are more pronounced under eyes-closed conditions, suggesting that sleep loss compromises sensory reweighting processes required to adapt to reduced sensory input. Similarly, acutely sleep-deprived individuals exhibit increased synchronization errors when adapting gait timing to a rhythmic auditory cue, indicating that supraspinal feedforward processes involved in movement timing and gait regulation are sensitive to sleep loss. Together, these findings suggest that motor impairments following sleep loss may arise not only from reduced cognitive resources, but also from disruptions in sensorimotor processing. However, the relative contribution of these mechanisms to impaired motor performance remains unclear.

Neuroimaging and neurostimulation evidence points to disruptions in both sensorimotor and attentional networks following sleep loss. Transcranial magnetic stimulation studies have shown that sleep deprivation reduces intracortical inhibition and facilitation in the primary motor cortex (11, 12), while somatosensory evoked potential studies have demonstrated increased excitability in the primary somatosensory cortex, suggesting that altered sensory gain may contribute to disrupted sensorimotor integration (13, 14). At the network level, resting-state fMRI studies have reported reduced functional connectivity between sensorimotor regions and subcortical structures (15), as well as decreased thalamocortical connectivity to somatosensory and motor cortex (16, 17). Sleep loss also consistently affects higher-order attentional networks, with reduced activation and connectivity in the dorsal attention network, particularly within the intraparietal sulcus and frontal eye fields (18–20). Critically, however, all of these studies have examined resting-state brain activity. Whether these network-level disruptions manifest during active motor task execution remains unknown.

To address this need, high-density electroencephalography (hdEEG) is well suited to simultaneously capture cortical dynamics across distributed sensorimotor and attentional networks during active motor tasks. By recording fluctuations in electrical potential generated by the collective activity of neurons through scalp electrodes, EEG enables real-time examination of brain activity with excellent temporal resolution and relatively good spatial resolution. Recent advances in artifact correction methods and the application of volume conduction models have further improved both signal quality during movement and spatial resolution, making it possible to distinguish activity across multiple cortical regions simultaneously (21–23). During movement, task-related cortical engagement is commonly indexed through event-related desynchronization (ERD), defined as a reduction in oscillatory power relative to a baseline period (24). Prior work suggests that changes in task demands, fatigue, and cognitive workload are accompanied by alterations in task-related desynchronization, reflecting shifts in cortical engagement during motor and cognitive performance (25–27). Yet, to date, no study has directly compared the contributions of sensorimotor and attentional network disruptions to motor impairments during active task performance under CSR.

The present study addressed this gap by investigating the effects of CSR on cortical oscillatory dynamics within sensorimotor and attentional networks during sustained submaximal ankle contractions. Ankle force production was selected as the motor task because it requires continuous sensorimotor feedback, precise force regulation, and sustained attention, making it sensitive to both sensorimotor and cognitive disruptions associated with sleep loss. Moreover, the ankle joint plays a central role in postural control and gait, and precise regulation of ankle force is fundamental to stable locomotion and fall prevention (28). EEG-derived ERD was used to index task-related cortical engagement across sensorimotor and attentional regions simultaneously during active motor performance. Based on the evidence reviewed, two mechanisms were hypothesized to contribute to motor impairments under CSR. First, consistent with the established sensitivity of attentional networks to sleep loss, altered ERD was expected within attentional control regions, reflecting diminished top-down cognitive engagement during force production. Second, given evidence that sleep loss disrupts sensorimotor integration and corticomotor excitability, altered ERD within sensorimotor cortex was also expected, independent of attentional network changes. Together, these hypotheses allowed for a direct assessment of the relative contributions of attentional and sensorimotor mechanisms to CSR-induced motor impairments.

## Methods

### Subjects

Twenty-four physically active male participants were initially recruited to take part in the study. Male participants were recruited exclusively to minimize variability attributable to menstrual cycle-related fluctuations in sleep architecture and neuromuscular performance (29, 30). Physical activity was defined in accordance with WHO guidelines as engaging in 150-300 minutes of moderate-intensity, or 75-150 minutes of vigorous-intensity aerobic physical activity per week, or an equivalent combination of both (31). Baseline sleep quality and daytime sleepiness were assessed using the Pittsburgh Sleep Quality Index (PSQI) and Epworth Sleepiness Scale (ESS), respectively (32, 33). Eligibility criteria required a PSQI score ≤5 and an ESS score <11 to ensure adequate habitual sleep quality and to exclude individuals with suspected sleep disturbances. Additional exclusion criteria included daily alcohol consumption, caffeine intake exceeding 400 mg/day, use of psychoactive substances, smoking, and the presence of metabolic, cardiovascular, respiratory, or orthopedic conditions that could interfere with sleep or motor performance.

Following data quality control, nine participants were excluded from the final analysis due to poor electroencephalographic signal quality (n = 7) or missing data (n = 2), resulting in a final sample of 15 participants. Participants had a mean age of 21.9 ± 2.1 years, height of 1.81 ± 0.08 m, and body mass of 75.9 ± 14.4 kg. Baseline sleep quality and daytime sleepiness were within normal ranges (PSQI: 3.3 ± 1.0; ESS: 3.7 ± 3.1). The study protocol was approved by the Ethics Committee of UZ Leuven (S67790) and conducted in accordance with the Declaration of Helsinki. Written informed consent was obtained from all participants before participation.

### Study design

Each participant completed two conditions in a randomized, counterbalanced crossover design: a chronic sleep restriction (CSR) condition and a control condition (CON, Fig. 1A). The conditions were separated by a one-week wash-out period. Prior to the first experimental session, participants underwent a familiarization session in which they were introduced to the PVT and motor tasks.

**Figure 1.**
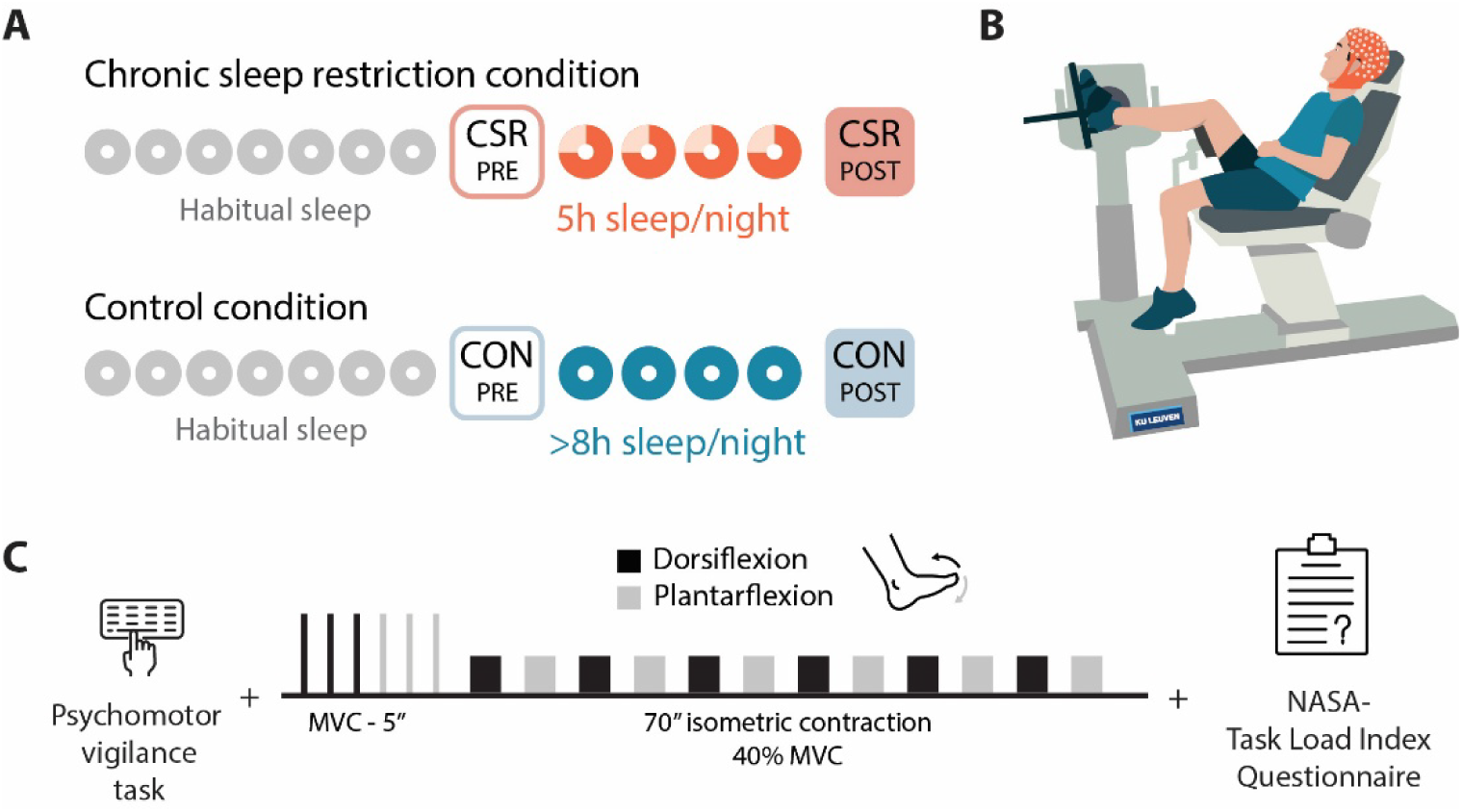
Overview of the experimental design and protocol. **(A)** Participants completed two conditions in a randomized counterbalanced crossover design: a chronic sleep restriction (CSR) condition, in which sleep was restricted to 5 hours per night for four consecutive nights, and a control (CON) condition, in which normal sleep habits (>8 hours per night) were maintained over the same period. Each condition comprised of a pre-intervention laboratory session (CSR_PRE_/CON_PRE_) and a post-intervention session (CSR_POST_/CON_POST_). Seven nights before each pre-intervention session, habitual sleep was monitored using actigraphy. **(B)** Participants were seated in an isokinetic testing chair with the dominant foot secured to the footplate, aligned at the lateral malleolus, to measure isometric ankle torque during dorsiflexion and plantarflexion. **(C)** Each laboratory session consisted of a 10-minute Psychomotor Vigilance Task (PVT), a motor task including six maximal voluntary contractions (MVCs, 5 s each) followed by twelve alternating submaximal isometric contractions at 40% MVC (70 s each) for both dorsiflexion and plantarflexion, and the NASA Task Load Index (NASA-TLX) questionnaire.

In both conditions, habitual sleep was monitored for seven consecutive nights prior to the first laboratory session (CSR_PRE_ or CON_PRE_). After this baseline session, participants entered the experimental manipulation phase. In the CSR condition, participants were instructed to restrict their sleep to 5 hours per night for four consecutive nights, a duration selected to mirror the length of a typical working week and to reflect the pattern of sleep restriction most commonly encountered in occupational and real-world settings. This was simulated using a late sleep restriction protocol in which bedtime remained habitual, but wake time was advanced. In the CON condition, participants were instructed to maintain their normal sleep habits over the same period. A minimum of 8 hours time in bed per night was encouraged. Following the four-night intervention period, all participants returned to the laboratory for a second experimental session (CSR_POST_ or CON_POST_).

Sleep duration and compliance were monitored throughout the study with wrist-worn ActiGraph wGT3X-BT accelerometers (ActiGraph LLC, Pensacola, FL, USA) and the consensus sleep diary (34, 35). Participants were excluded from analysis if their total sleep time, as recorded in the sleep diary, deviated by more than 60 minutes from the prescribed sleep schedule on more than one night during the sleep restriction period. Participants were instructed to maintain similar dietary intake across conditions, to abstain from alcohol and caffeine for 24 hours prior to each experimental session, and to refrain from napping during the study period. They were also asked to avoid intense physical activity during this period. All experimental sessions were conducted in the afternoon, with testing starting between 13:00 and 15:00. Importantly, session timing was kept consistent within each participant across both conditions to control for potential circadian effects.

### PVT

Sustained attention was measured using a 10-minute Psychomotor Vigilance Task (PVT). The PVT is a well-established and highly sensitive measure of sleep loss and was included as a manipulation check to confirm the effectiveness of the sleep restriction protocol (36). Participants were instructed to respond as quickly as possible to visual stimuli presented at random inter-stimulus intervals on a computer screen. Both reaction times and lapses (responses >500 ms) were recorded.

### Motor Task

Participants were seated in an isokinetic dynamometer (Biodex System 3, Biodex Medical Systems, Shirley, NY, USA) chair with the backrest reclined at 85 degrees (Fig. 1B). The hips were flexed at 120 degrees, resulting in a more reclined pelvis and a slightly posterior trunk tilt rather than a fully upright position. The thighs were angled downward toward the seat, and the dominant leg was elevated and extended forward, supported by the dynamometer arm, positioning the knee in slight flexion (160 degrees). The foot was placed on the footplate of the dynamometer, with the lateral malleolus aligned with the rotational axis of the device to ensure accurate torque measurement. Body and limb positioning were carefully adjusted during the familiarization session and replicated across all experimental sessions to ensure consistency. The lower leg and foot were securely fastened using rigid straps to minimize movement and isolate ankle joint torque production. The trunk was further stabilized with a shoulder harness, and additional straps across the chest and pelvis were applied to reduce compensatory movements during maximal force production tasks.

Before the motor task protocol, participants performed submaximal plantarflexion (PF) and dorsiflexion (DF) contractions of progressively increasing intensity as a familiarization and warm-up procedure. Subsequently, the protocol commenced with three maximal voluntary contractions (MVCs) for both DF and PF, each lasting 5 s and separated by a 1-minute rest interval. Following the MVC assessments, participants performed twelve submaximal contractions at 40% of their MVC torque, alternating between dorsiflexion and plantarflexion (Fig. 1C). During these trials, visual feedback of the real-time torque signal was provided on a computer screen positioned in front of the participants. Subjects were instructed to maintain torque output as close as possible to a horizontal target line displayed on the screen. Each submaximal contraction lasted 70 seconds and was followed by a 30-second rest period. Throughout the complete protocol, no verbal encouragement was provided.

### NASA Task Load Index (NASA-TLX)

After completing the testing protocol, participants rated their perceived workload using a modified version of the NASA Task Load Index (NASA-TLX; (37)). This questionnaire was used to assess workload experienced during the complete motor task and comprised subscales for mental demand, physical demand, frustration, performance, and effort.

### Instrumentation and data acquisition

#### Dynamometry

The raw torque signal was scaled, rectified, and low-pass filtered using a fourth-order Butterworth filter with a 10 Hz cutoff frequency. For each session, maximal torque was determined as the highest value recorded across the three MVC trials. Target torque for submaximal contractions was set at 40% of the session-specific MVC. For submaximal contractions, the middle 60 seconds of each trial (seconds 5 to 65) were used for analysis to minimise the influence of task initiation and termination effects and better capture steady-state performance. Torque variability was quantified as the coefficient of variation (CV, standard deviation/mean), and tracking accuracy was assessed as the root mean square error (RMSE) relative to the target torque.

#### Electroencephalography

Brain activity was recorded using a 128-channel high-density EEG system (hdEEG; ActiCHamp amplifier, Brain Products GmbH, Gilching, Germany) at a sampling rate of 1,000 Hz, with electrode FCz as the online recording reference. Data acquisition during the motor tasks was performed using BrainVision Recorder (Brain Products GmbH). EEG data were analyzed using the Noninvasive Electrophysiology Toolbox (NET; (38)), an open-source MATLAB-based toolbox for large-scale automated hdEEG analysis. The analysis pipeline comprised four sequential stages: head modelling, signal preprocessing, source reconstruction, and event-related spectral analysis.

### Head modelling

Since individual structural MRI data were not available, a template-based head model was constructed. The Brain Products 128-channel template electrode positions were rigidly co-registered to a standard Montreal Neurological Institute (MNI) template MRI. Head tissue segmentation was performed by warping a pre-segmented template image to the template MRI, thus identifying 12 tissue compartments (39). Conductivity values for each tissue compartment were assigned based on predefined values from the literature (40). The leadfield matrix, describing the linear relationship between scalp EEG signals and underlying neural sources, was computed using the Generalized Finite Difference Method (GFDM) with a dipole grid resolution of 6 mm (41).

### Signal preprocessing

Bad channels were identified automatically based on signal correlation with neighboring channels and noise variance, using an outlier threshold of mean + 3 standard deviations. Detected bad channels were reconstructed through spatial interpolation of surrounding electrodes, to maintain spatial accuracy. The continuous EEG signal was then band-pass filtered between 1 and 80 Hz, followed by downsampling from 1,000 to 200 Hz to reduce computational load while retaining signal content and variance. Signal despiking was applied after artifact attenuation.

Four categories of biological artifacts were removed sequentially using blind source separation (BSS; (21)). Ocular artifacts were identified using a FastICA deflation algorithm, with components detected based on kurtosis (threshold: 12). Movement artifacts were attenuated using a symmetric FastICA algorithm, with detection based on sample entropy (threshold: 0.8), applied over 30-second windows with a 7 Hz low-pass filter. Myogenic artifacts were removed using Independent Vector Analysis (IVA), with components identified based on the ratio of gamma-band to total-band power (threshold: 0.5). Cardiac artifacts were identified using a FastICA deflation algorithm based on signal skewness (threshold: 1). For each artifact type, artifactual independent components were automatically identified and subtracted, after which artifact-free data were reconstructed. Finally, signals were re-referenced to the common average.

### Source reconstruction

Neural source activity was estimated from the preprocessed EEG data using exact low-resolution electromagnetic tomography (eLORETA; (42)), with a regularization parameter lambda of 0.05. A spatial filter combining the leadfield matrix and the preprocessed signals was applied to reconstruct neural activity time courses at each voxel of the template brain, with source maps registered to MNI space at 4 mm resolution with 8 mm smoothing.

### Event-related spectral analysis

Event-related synchronization and desynchronization (ERS/ERD) were computed to quantify task-related modulations of oscillatory power across four frequency bands: theta (4-8 Hz), alpha (8-13 Hz), beta (13-30 Hz), and gamma (30-50 Hz). A short-time Fourier transform was applied to the source-reconstructed signals to obtain time-frequency decompositions. Epochs were extracted from -10 s to +80 s relative to contraction onset, with contraction offset occurring at +70 s. Power changes were expressed as relative change with respect to a pre-contraction baseline period of -10 s to -2 s, thereby excluding the 2 s immediately preceding contraction onset to avoid contamination from anticipatory neural activity.

ERS/ERD were analyzed separately for dorsiflexion and plantarflexion contractions and extracted for two large-scale brain networks: the sensorimotor network (SMN) and the dorsal attention network (DAN). Based on previous literature, regions of interest (ROIs) for the SMN included the primary motor cortex (M1), the primary somatosensory cortex (S1), and the supplementary motor area (SMA); ROIs for the DAN included the intraparietal sulcus (IPS), the superior parietal lobule (SPL), and the frontal eye fields (FEF; see Table 1 for MNI coordinates). For M1 and S1, only the dominant hemisphere seed was included, given the lateralized nature of unilateral ankle force production. Foot dominance was determined via self-report, with participants asked to identify the foot they would naturally use to shoot a football. For all other ROIs, left and right hemisphere seeds were averaged. Network-level ERS/ERD estimates were then obtained by averaging across all constituent ROIs within each network.

**Table 1.** MNI coordinates of regions of interest for the sensorimotor and dorsal attention networks.

| Region | Hemisphere | MNI coordinates (x, y, z) |
| --- | --- | --- |
| Sensorimotor network (SMN) |  |  |
| Primary motor cortex (M1) | Left | -34, -24, 60 |
| Primary motor cortex (M1) | Right | 36, -22, 58 |
| Supplementary motor area (SMA) | Bilateral | 0, -8, 56 |
| Primary somatosensory cortex (S1) | Left | -56, -20, 18 |
| Primary somatosensory cortex (S1) | Right | 56, -20, 18 |
| Dorsal attention network (DAN) |  |  |
| Intraparietal sulcus (IPS) | Left | -34, -48, 44 |
| Intraparietal sulcus (IPS) | Right | 32, -56, 44 |
| Superior parietal lobule (SPL) | Left | -12, -60, 56 |
| Superior parietal lobule (SPL) | Right | 30, -62, 62 |
| Frontal eye field (FEF) | Left | -35, -5, 52 |
| Frontal eye field (FEF) | Right | 35, -5, 52 |

#### Actigraphy

Actigraphy data were processed and analyzed using the GGIR package in R (version 3.2-6), which performs autocalibration, detects non-wear time, and extracts sleep and physical activity metrics based on raw acceleration signals (43).

### Statistical analysis

All statistical analyses were performed in R (version 2025.05.1) using the lme4 and lmerTest packages. Model assumptions were assessed through visual inspection of residual and Q-Q plots to evaluate homoscedasticity and normality of residuals. Sleep duration and sleep efficiency were assessed using actigraphy and the consensus sleep diary. For each participant, nightly values were averaged across the intervention nights per condition, and the resulting means were entered into a linear mixed model (LMM) with condition (CSR, CON) and session (PRE, POST) as fixed effects and participant as a random intercept. All LMMs included condition order (CSR-first vs. CON-first) as an additional covariate to account for potential carryover effects inherent to the crossover design.

PVT outcomes (median reaction time, lapses) were analyzed using LMMs with condition and session as fixed effects and participant as a random intercept. For maximal voluntary contraction (MVC) torque, the peak torque value across the three MVC trials was selected per participant per session per movement direction (dorsiflexion and plantarflexion separately), and analyzed using an LMM with condition, session, movement direction, and their three-way interaction as fixed effects and participant as a random intercept. For submaximal contractions, torque variability (CV) and tracking error (RMSE) were analyzed using LMMs with condition, session, movement direction (dorsiflexion, plantarflexion), trial (1–6), and their interactions as fixed effects and participant as a random intercept. Trial was included as a fixed factor to account for potential within-session changes in performance (e.g., muscle fatigue or adaptation across repeated contractions), and movement direction was included to test whether condition and session effects differed between dorsiflexion and plantarflexion. Models were fitted with REML = FALSE.

For brain oscillatory activity, the mean ERS/ERD (%) per trial was extracted for each frequency band (theta, alpha, beta, gamma) and network (sensorimotor, dorsal attention), averaged across all ROIs within each network. A single LMM was fitted with condition, session, frequency band, movement direction, network, and trial number as fixed effects, along with condition order as a covariate and participant as a random intercept. Rather than fitting separate models per movement condition, network, and frequency band, three targeted three-way interactions were included to address four pre-specified research questions: whether the overall condition x session effect was significant (Q1), and whether it was moderated by network (Q2), movement direction (Q3), or frequency band (Q4). More complex random-effects structures (e.g., random slopes for session or condition) were explored but not retained because of convergence issues and/or singular fits, and because they did not improve model fit. The four omnibus interaction tests (Q1-Q4) were not corrected for multiple comparisons as they address distinct research questions. Where omnibus tests were significant or where exploratory follow-up was warranted, post hoc interaction contrasts were computed using emmeans, with FDR correction (Benjamini-Hochberg) applied separately within each family of contrasts (by frequency band, movement direction, and network) (44).

NASA Task Load Index subscales (mental demand, physical demand, performance, effort, and frustration) and the total score were each analyzed using LMMs with condition and session as fixed effects and participant as a random intercept. Where a significant condition by session interaction was observed, within-condition PRE vs. POST contrasts were performed using emmeans. For non-EEG outcomes, no correction for multiple comparisons was applied given the exploratory nature of these analyses and the limited number of models per outcome domain; the significance threshold was set at α = .05. For EEG outcomes, FDR-corrected p-values were used as described above.

## Results

### Sleep Assessment

To verify the effectiveness of the sleep restriction manipulation, total sleep duration was analyzed using actigraphy as well as the consensus sleep diary (Fig. 2). For the actigraphy, a significant condition × session interaction was observed, *F*(1,45) = 56.09, *p* < .001. Within-condition comparisons further showed a significant reduction in sleep duration from session CSR_PRE_ to session CSR_POST_ in the experimental condition (Δ = - 2.36 h, *SE* = 0.27, *p* < .001), while no significant change was observed in the control condition (*p* = .188). Sleep efficiency showed a significant condition × session interaction, *F*(1, 45) = 5.20, *p* = .027, driven by a small increase in the experimental condition from PRE (83.6% ± 4.6%) to POST (87.7% ± 3.9%; Δ = +4.1%, *p* = .009), while no change was observed in the control condition (PRE: 86.0% ± 4.5%; POST: 85.4% ± 6.9%; *p* = .68).

**Figure 2.**
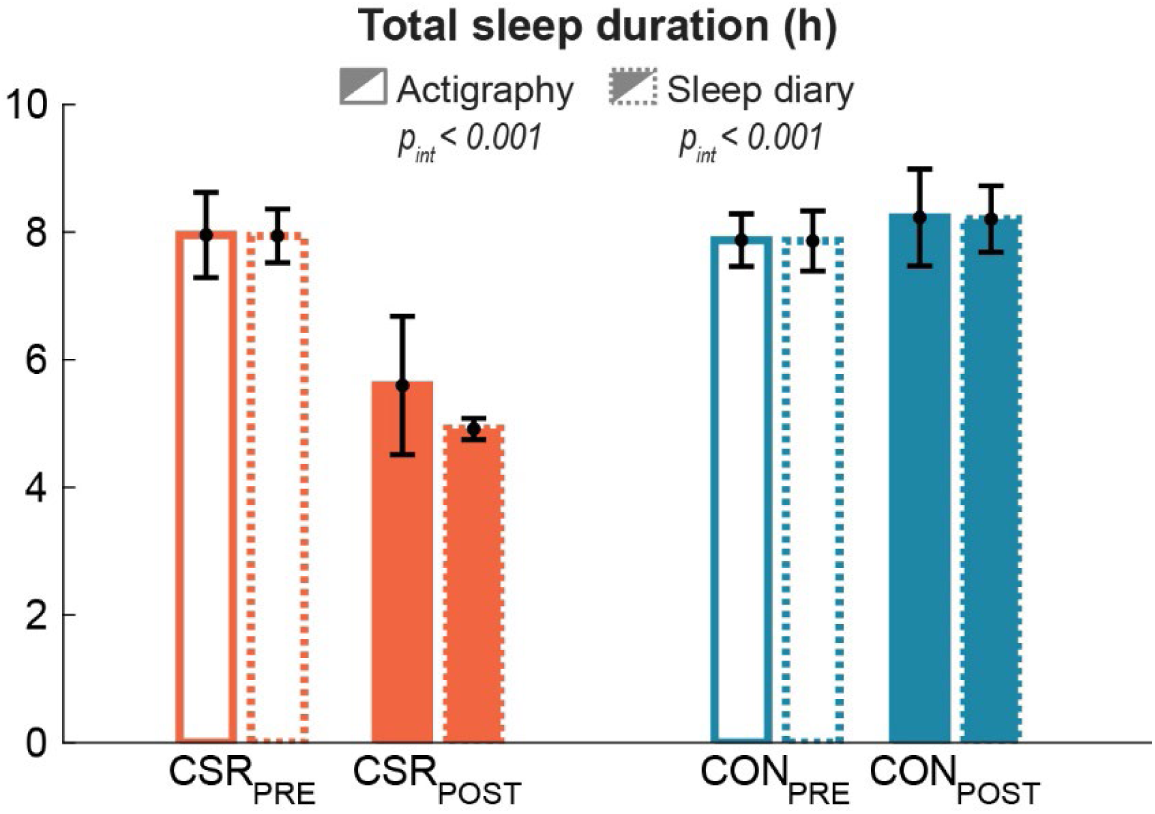
Total sleep duration (h) measured via actigraphy (solid bars) and sleep diary (dashed bars) across sessions for the chronic sleep restriction (CSR) and control (CON) conditions. Both measurement methods showed a significant reduction in sleep duration from CSR_PRE_ to CSR_POST_, confirming the effectiveness of the sleep restriction manipulation. P_int_ = condition × session interaction effect.

For the sleep diary data, a consistent pattern was observed, with a significant condition × session interaction, *F*(1, 42) = 153.12, *p* < .001. Within-condition comparisons further showed a significant reduction in sleep duration from PRE to POST in CSR (Δ = −3.03 h, *p* < .001).

### Psychomotor Vigilance Task

Psychomotor vigilance performance was assessed using median reaction time and the number of lapses (reaction times > 500 ms). **Median reaction time** showed a significant condition × session interaction (F(1, 45) = 8.46, p = .006, Fig. 3A). Post-hoc within-condition comparisons revealed that median reaction time increased significantly from PRE to POST in the Experimental (CSR) condition (Δ = +30.97 ms, SE = 9.28, p = .002), indicating a slowing of responses following sleep restriction. No significant change was observed in the Control condition from PRE to POST (Δ = −5.93 ms, SE = 9.28, p = .526).

**Figure 3.**
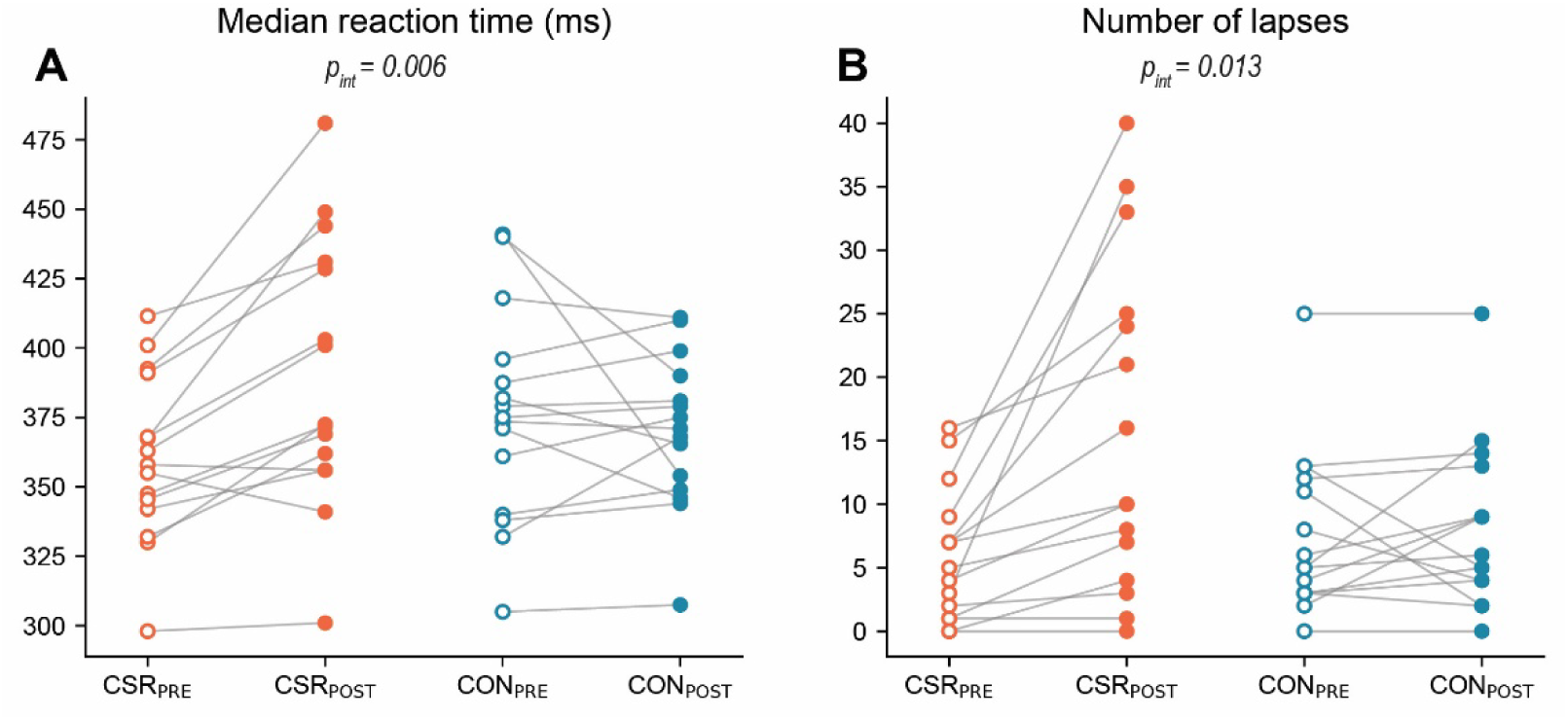
Individual data points for **(A)** median reaction time (ms) and **(B)** number of lapses (reaction times > 500 ms) across sessions for the chronic sleep restriction (CSR, orange) and control (CON, blue) conditions. Open circles represent PRE values and filled circles represent POST values, with grey lines connecting each individual’s PRE and POST scores. A significant condition × session interaction (P_int_) was observed for both median reaction time and number of lapses.

**Lapses** also showed a significant condition × session interaction (F(1, 45) = 6.67, p = .013, Fig. 3B). Within-condition post-hoc comparisons revealed a significant increase in the number of lapses from PRE to POST in the Experimental condition (Δ = +9.87, SE = 2.63, p < .001), while no significant change was observed in the Control condition (Δ = +0.60, SE = 2.63, p = .820). Together, these results indicate that chronic sleep restriction significantly impaired sustained attention, as reflected by both slower reaction times and a greater number of attentional lapses following the restriction period, with no such changes occurring under normal sleep conditions.

### Motor Task

**Maximal torque** did not show a significant condition × session interaction (F(1, 105) = 0.00, p = .958, Fig. 4A and 4D), indicating that changes in force production over time were comparable between the CSR and control conditions. There was a strong main effect of movement direction (F(1, 105) = 1177.33, p < .001), reflecting substantially higher torque values during plantarflexion compared with dorsiflexion. The condition × session × movement direction interaction was also not significant (F(1, 105) = 0.00, p = .947), indicating that the absence of CSR-related effects over time was consistent across both movement directions.

**Figure 4.**
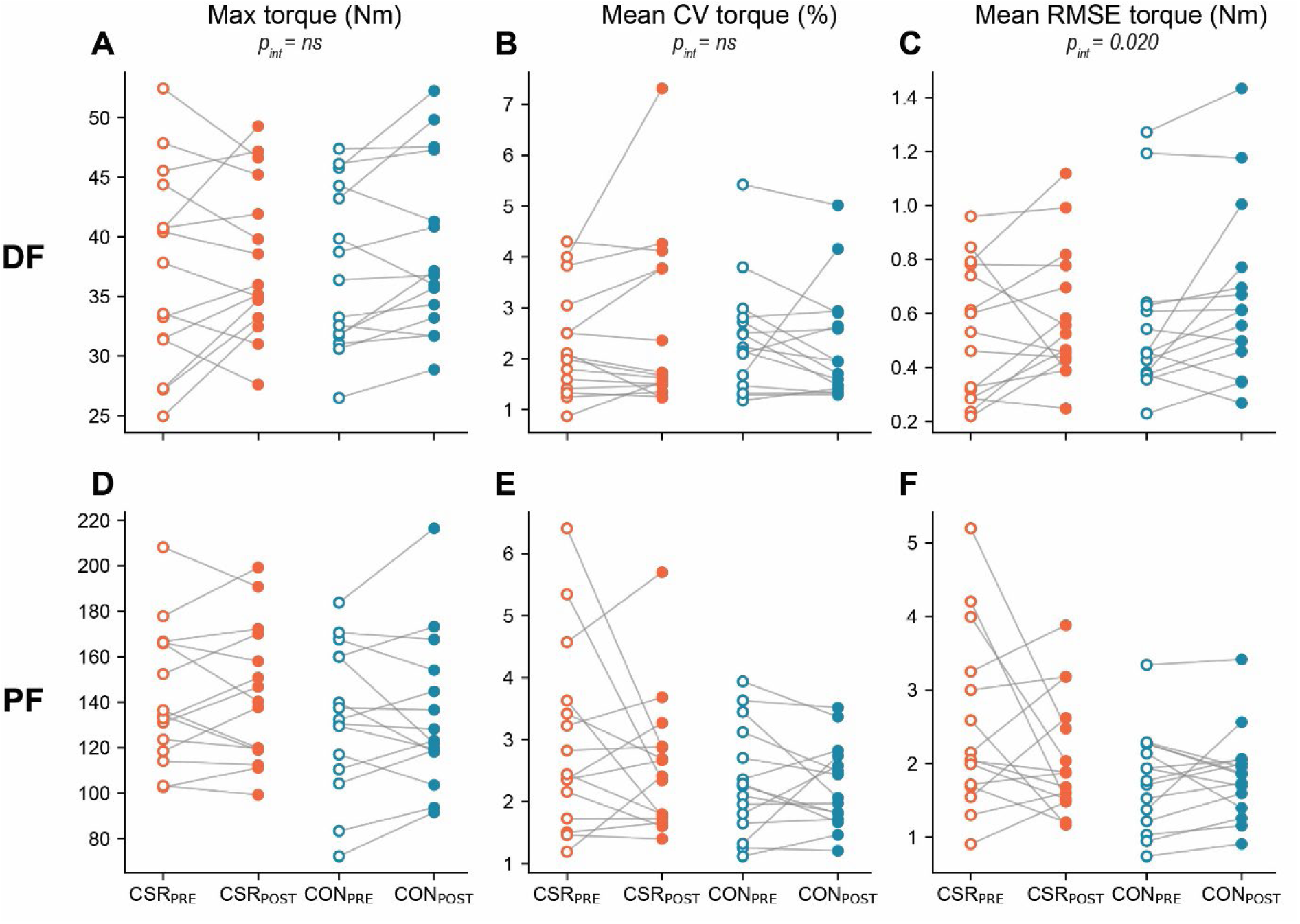
Individual data points for maximal and submaximal torque outcomes during dorsiflexion (DF; panels A-C) and plantarflexion (PF; panels D-F) across sessions for the chronic sleep restriction (CSR, orange) and control (CON, blue) conditions. Open circles represent PRE values and filled circles represent POST values, with grey lines connecting each individual’s PRE and POST scores. Panels A and D show maximal torque (Nm); panels B and E show torque variability during submaximal contractions expressed as the coefficient of variation (CV, %); panels C and F show tracking error during submaximal contractions expressed as root mean square error (RMSE, Nm). P_int_ = condition × session interaction.

For submaximal contractions, **torque variability (CV)** showed no significant condition × session interaction (F(1, 703) = 0.03, p = .864). In addition, there was no evidence that this effect differed by contraction type (condition × session × contraction type: F(1, 703) = 2.91, p = .089, Fig. 4B and 4E). For **torque tracking error (RMSE)**, a significant condition × session interaction was observed (F(1, 703) = 5.40, p = .020; Fig. 4C and 4F), together with a trend towards a significant condition × session × contraction type interaction (F(1, 703) = 3.67, p = .056), indicating that the CSR-related change over time depended on movement direction. This was further supported by a strong main effect of contraction type (F(1, 703) = 626.16, p < .001), with RMSE for plantarflexion being higher than for dorsiflexion. Exploratory post-hoc comparisons were conducted to decompose the interaction structure. These revealed a condition × session interaction for plantarflexion (p = .003), but not for dorsiflexion (p = .776). This was driven by a decrease in RMSE from PRE to POST in the experimental condition (p < .001), with no corresponding change in the control condition. Together, these results indicate that torque tracking accuracy during plantarflexion showed an unexpected improvement from PRE to POST specifically in the CSR condition. The basis for this finding is unclear and was not predicted a priori.

### Brain Activity

To assess the effects of CSR on cortical oscillatory activity during sustained isometric contractions, we fitted a single linear mixed model with a random intercept per subject. The model first tested the overall condition x session effect collapsed across all frequency bands, movement directions, and networks (Q1), and then examined whether this effect was moderated by network (Q2), movement direction (Q3), or frequency band (Q4). Omnibus F-tests for each question are reported in Table 2; post hoc interaction contrasts are reported in Table 3.

**Table 2.** Omnibus tests for the condition x session interaction and its moderation by frequency band, movement direction, and network.

| Main Omnibus Test Results | F(df1, df2) | p |
| --- | --- | --- |
| Condition x session (Q1) | F(1, 5713) = 14.20 | < .001 |
| Condition x session x network (Q2) | F(1, 5713) = 0.17 | .682 |
| Condition x session x movement direction (Q3) | F(1, 5713) = 9.13 | .003 |
| Condition x session x frequency band (Q4) | F(3, 5713) = 1.80 | .146 |

**Table 3.** Post hoc interaction contrasts for the condition x session effect on ERS/ERD by frequency band, movement direction, and network.

| Post hoc test results |  | Estimate | SE | pFDR |
| --- | --- | --- | --- | --- |
| Post hoc: Overall sleep effect (Q1) | CSR: PRE vs POST | 3.60 | 0.85 | < .001 |
|  | Control: PRE vs POST | -0.93 | 0.85 | .276 |
|  | Difference (CSR vs Control) | -4.52 | 1.20 | < .001 |
| Post hoc: By network (Q2) | Sensorimotor network | -5.02 | 1.70 | .006 |
|  | Attention network | -4.03 | 1.70 | .018 |
| Post hoc: By movement direction (Q3) | Dorsiflexion | -8.15 | 1.70 | < .001 |
|  | Plantarflexion | -0.90 | 1.70 | .598 |
| Post hoc: By frequency band (Q4) | Theta | -0.06 | 2.40 | .981 |
|  | Alpha | -5.57 | 2.40 | .041 |
|  | Beta | -4.79 | 2.40 | .061 |
|  | Gamma | -7.68 | 2.40 | .006 |

The **overall condition x session** interaction (**Q1**) was significant (F(1, 5713) = 14.20, p < .001), indicating that the PRE-to-POST change in ERS/ERD differed between the CSR and control conditions across all bands, directions, and networks. Post hoc contrasts confirmed a significant PRE-to-POST increase in ERD in the CSR condition (estimate = 3.60, SE = 0.85, p < .001) but not in the control condition (estimate = -0.93, SE = 0.85, p = .276), with the between-condition difference in PRE-to-POST change estimated at 4.52% (SE = 1.20, p < .001).

The condition x session x **network interaction (Q2)** was not significant (F(1, 5713) = 0.17, p = .682), indicating that the sleep-related change in ERD did not differ between the sensorimotor and attention networks. Exploratory post hoc contrasts confirmed that the differential condition x session effect was present in both networks (SMN: estimate = -5.02, SE = 1.70, pFDR = .006; DAN: estimate = -4.03, SE = 1.70, pFDR = .018).

The condition x session x **movement direction** interaction (**Q3**) was significant (F(1, 5713) = 9.13, p = .003), indicating that the sleep-related increase in ERD was specific to dorsiflexion. Post hoc contrasts confirmed a significant differential effect during DF (estimate = -8.15, SE = 1.70, pFDR < .001) and no effect during PF (estimate = -0.90, SE = 1.70, pFDR = .598; Figure 5 vs. Figure 6).

**Figure 5.**
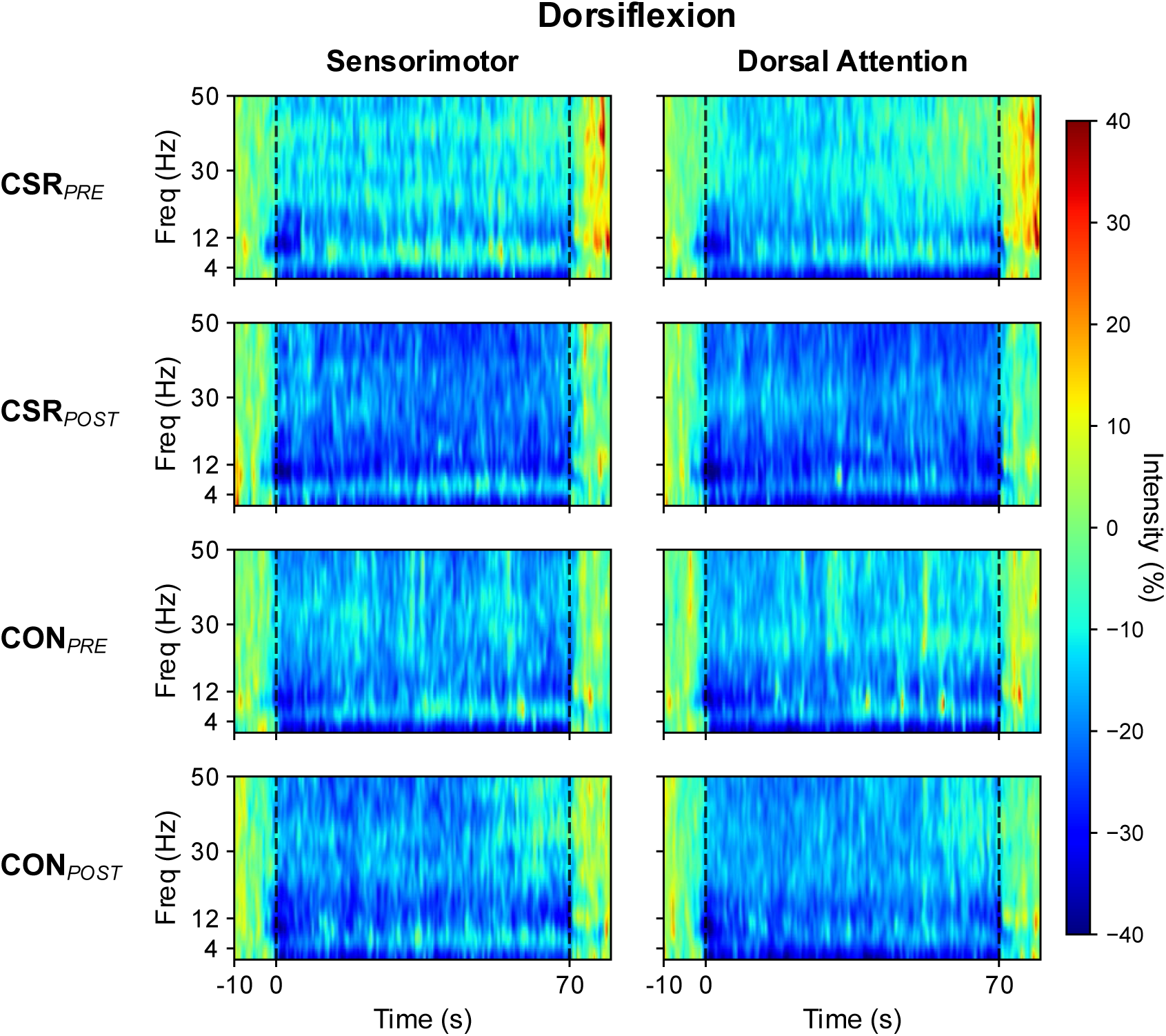
Time–frequency representations of ERS/ERD during sustained dorsiflexion for the sensorimotor and dorsal attention networks across sleep conditions and sessions. Mean oscillatory activity (%) is shown from 1–50 Hz for the chronic sleep restriction (CSR) and control (CON) conditions at PRE and POST assessments. Dashed vertical lines indicate task onset (0 s) and offset (70 s). Negative values (blue) reflect event-related desynchronization (ERD) and positive values (yellow/red) reflect event-related synchronization (ERS).

**Figure 6.**
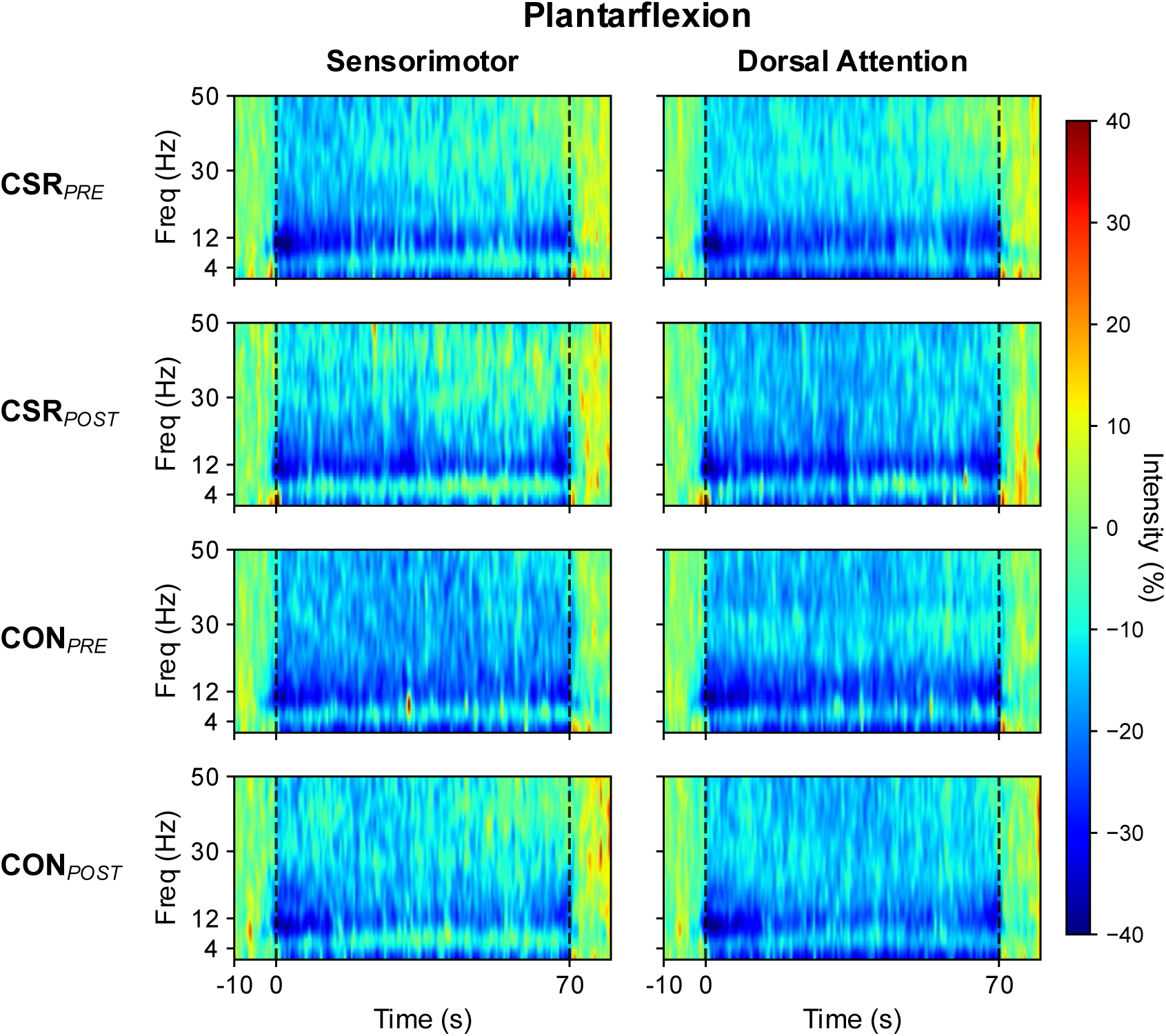
Time–frequency representations of ERS/ERD during sustained plantarflexion for the sensorimotor and dorsal attention networks across sleep conditions and sessions. Mean oscillatory activity (%) is shown from 1–50 Hz for the chronic sleep restriction (CSR) and control (CON) conditions at PRE and POST assessments. Dashed vertical lines indicate task onset (0 s) and offset (70 s). Negative values (blue) reflect event-related desynchronization (ERD) and positive values (yellow/red) reflect event-related synchronization (ERS).

Finally, regarding **frequency band** specificity (**Q4**), the condition x session x frequency band interaction was not significant (F(3, 5713) = 1.80, p = .146), indicating that the sleep-related increase in ERD was not formally confined to a specific frequency band. Nonetheless, exploratory analyses of interaction contrasts by band revealed that the differential condition x session effect was largest and most reliable in the gamma band (estimate = -7.68, SE = 2.40, pFDR = .006) and the alpha band (estimate = -5.57, SE = 2.40, pFDR = .041), with beta showing a trend in the same direction (pFDR = .061) and theta showing no effect (pFDR = .981). However, the absence of a significant omnibus interaction means frequency band specificity cannot be formally claimed.

A significant linear effect of trial number was observed (F(1, 5713) = 15.91, p < .001), reflecting a progressive increase in ERD across successive trials within a session. Condition order did not significantly affect ERS/ERD (F(1, 13) = 0.12, p = .735), suggesting no systematic carryover effects between the CSR and control conditions.

### NASA Task Load Index

NASA Task Load Index scores are presented in Table 4. A significant condition by session interaction was observed for mental demand (p = 0.027). Post hoc contrasts revealed that mental demand increased significantly from PRE to POST in the CSR condition (estimate = 4.5, SE = 1.24, t(48) = 3.67, p < 0.001), whereas no significant change was observed in the control condition (p = 0.592). No significant condition by session interactions were found for physical demand, performance, effort, frustration, or total score (all p > 0.05).

**Table 4.** Descriptive statistics (mean ± std) for the NASA-TLX subscales and p-value for condition × session interaction.

| Outcome | CSR<sub>PRE</sub> | CSR<sub>POST</sub> | CON<sub>PRE</sub> | CON<sub>POST</sub> | p |
| --- | --- | --- | --- | --- | --- |
| Mental demand | 6.7 ± 4.5 | 11.3 ± 6.1 | 7.2 ± 4.4 | 7.9 ± 4.7 | **0.027** |
| Physical demand | 15.5 ± 1.7 | 15.9 ± 3.5 | 13.9 ± 2.0 | 13.9 ± 2.3 | 0.755 |
| Performance | 5.7 ± 3.2 | 7.3 ± 3.3 | 6.4 ± 3.9 | 5.9 ± 4.3 | 0.108 |
| Effort | 15.4 ± 2.8 | 15.4 ± 4.2 | 14.9 ± 2.1 | 14.1 ± 2.6 | 0.508 |
| Frustration | 6.1 ± 5.1 | 6.9 ± 4.9 | 3.8 ± 3.0 | 4.9 ± 4.1 | 0.870 |
| Total score | 49.4 ± 8.6 | 56.8 ± 11.5 | 46.3 ± 7.9 | 46.8 ± 8.2 | 0.086 |

## Discussion

The present study investigated the effects of chronic sleep restriction on cortical oscillatory dynamics during sustained isometric ankle contractions using high-density EEG source imaging. Following CSR, ERD increased relative to the control condition, and this effect was present across both the sensorimotor and dorsal attention networks. The increase in ERD was specific to dorsiflexion and was absent during plantarflexion. The effect was not confined to a specific frequency band. These neural changes were accompanied by increased subjective mental demand and a deterioration of sustained attention on the PVT, while objective motor performance was largely unimpaired. The observed increase in ERD following CSR without motor performance changes may reflect a compensatory upregulation of cortical resource allocation, whereby additional neural engagement across sensorimotor and attentional networks was required to maintain motor output under sleep pressure.

### Motor performance and compensatory cortical recruitment

No significant impairments in maximal strength or submaximal force control were observed following CSR. This is consistent with the broader literature, where effects of sleep loss on isometric force production and force steadiness are inconsistent and task-dependent. Some studies report impaired grip and load force control following sleep deprivation (45, 46), while others find no change or even reduced torque fluctuations during sustained knee contractions (47). The force control task used here involved a single-joint isometric contraction with continuous visual feedback, conditions that constrain motor complexity and enable real-time error correction. This could have potentially masked subtle CSR-induced impairments that might emerge in more dynamic or multijoint tasks. Importantly, preserved behavioral output does not necessarily imply that CSR was without effect: it is well established that individuals can maintain performance under sleep pressure by increasing cognitive effort, at the cost of greater subjective load (48, 49). Consistent with this, subjective mental demand was elevated under CSR in the present study despite unchanged force output, suggesting the task became more costly to sustain. The unexpected reduction in plantarflexion RMSE following CSR, while not readily explained by the experimental manipulation or condition order, does not alter this interpretation. No evidence of CSR-induced performance decrements was found, and the overall pattern across outcomes remains consistent with maintained motor output.

At the neural level, this dissociation between preserved performance and increased effort cost is mirrored by a broadband increase in ERD following CSR. ERD reflects a reduction in oscillatory power relative to a resting baseline and is the canonical marker of task-related cortical engagement (50): greater ERD indicates stronger or more widespread active cortical processing during the task. The observation that cortical engagement increased under CSR while motor output remained stable is therefore consistent with a compensatory account, in which the brain mobilizes additional neural resources to close the gap between reduced capacity and required output. This pattern has been documented across both sleep loss and motor fatigue literature. Drummond et al. showed using fMRI that total sleep deprivation was associated with increased prefrontal and parietal activation during cognitive tasks, scaling with task difficulty (51–53), and similar findings have been reported under CSR specifically (54, 55). In the motor domain, fatigue-related redistribution of cortical oscillatory activity has similarly been interpreted as compensatory support for declining motor drive (56, 57). Taken together, the co-occurrence of preserved force output, elevated subjective demand, and increased ERD in the present study is consistent with a scenario in which participants maintained performance by recruiting additional cortical resources.

However, this interpretation carries an implicit causal claim that the present data cannot directly test. It assumes that the increased cortical engagement was functionally necessary to maintain output, whereas it may equally reflect a concurrent neural response to sleep pressure that was not strictly required at this force level and degree of task familiarity. Therefore, an alternative interpretation is that the increased ERD reflects disrupted rather than augmented cortical engagement, consistent with sleep-related reductions in sustained attention and attentional network activity (20, 58). However, several considerations argue against this account. The direction of the ERD change, reflecting greater cortical desynchronization following CSR, is difficult to reconcile with genuine disengagement, which would more plausibly manifest as reduced rather than increased task-related activity. Motor performance was preserved, weakening the case for functionally meaningful attentional withdrawal during the task. Finally, no direct measure of within-task attention was obtained. Thus, the PVT impairment, while consistent with reduced vigilance capacity, cannot establish that attentional monitoring of force output was compromised during the contraction itself. We therefore regard attentional disengagement as an interpretation the present data do not support well, while acknowledging it cannot be fully excluded without a task-embedded attentional measure.

### Parallel changes across sensorimotor and dorsal attention networks

The hypotheses entering this study anticipated potentially separable contributions from the sensorimotor and dorsal attention networks. However, the ERD increase following CSR was observed simultaneously in both networks. The absence of a significant difference between networks is informative: it suggests that CSR did not selectively impair one system over the other during active motor performance. When both networks are actively co-recruited to support the same task, sleep pressure may affect them as a functional unit rather than selectively targeting one.

This co-engagement is not unexpected from a task demands perspective. Sustained force production at 40% MVC with continuous visual feedback simultaneously demands sensorimotor integration and top-down attentional monitoring of the torque signal. The DAN in particular supports precisely this kind of visuospatial, target-directed behavior (59). EEG research on isometric force maintenance has shown that activity during such tasks reflects the combined recruitment of sensorimotor and attentional networks, and that coupling between these networks is directly associated with force control performance (60). Rowe et al. further demonstrated that directing attention to a motor task specifically enhanced effective connectivity between prefrontal and premotor regions, indicating that attentional and motor systems operate in a coordinated manner during monitored motor performance (61). The parallel ERD increase across both networks following CSR may therefore reflect a disruption of this ordinarily coordinated recruitment rather than two independent processes responding separately to sleep pressure.

However, an important alternative deserves acknowledgment: the parallel change across SMN and DAN may not reflect a network-specific effect at all, but rather a more global shift in cortical state induced by CSR. Because the present study analyzed only two predefined networks, it is not possible to determine whether the ERD increase was selective to these networks or whether it would have appeared in any cortical network examined. Evidence from resting-state fMRI indicates that sleep deprivation induces widespread alterations in large-scale functional connectivity spanning multiple intrinsic brain networks, including the default mode, dorsal attention, frontoparietal, somatomotor, and thalamocortical systems (62–64). These findings raise the possibility that the SMN and DAN changes observed here reflect a broader cortical response to increasing sleep pressure rather than a network-specific effect. This limits the strength of conclusions about coordinated SMN-DAN recruitment specifically, and future studies including a broader set of networks or whole-brain analyses would be needed to determine whether the effect is genuinely network-selective.

Finally, it should be noted that the present analyses were based on power changes within predefined network seeds, and statements about network coupling or coordinated processing remain hypothetical. Power changes tell you that each network became more active, but not whether the two networks became more coordinated with each other. Compensation through network-level reorganisation, as described in the sleep deprivation neuroimaging literature (51, 52), implies increased inter-network coupling rather than just parallel local changes. Without connectivity analyses, the parallel ERD increase is consistent with coordinated recruitment but does not demonstrate it, and this distinction should be kept in mind when interpreting the network-level findings.

### Movement specificity: dorsiflexion vs. plantarflexion

The ERD changes were specific to dorsiflexion and absent during plantarflexion. This asymmetry was not predicted a priori but can be interpreted in light of known differences in corticospinal control between these movements. Brouwer and Ashby showed that motor cortex stimulation produces stronger short-latency facilitation of tibialis anterior motoneurons than plantarflexors, reflecting a more direct corticospinal influence on dorsiflexion muscles (65, 66). Lauber et al. similarly reported greater corticospinal excitability and stronger task-dependent modulation of intracortical inhibition in tibialis anterior compared to soleus (67). Neuroimaging studies also show that dorsiflexion engages broader cortical networks, including primary motor cortex and supplementary motor areas, than plantarflexion (68). Together, these findings suggest that dorsiflexion relies more heavily on cortical control, whereas plantarflexion depends relatively more on spinal and subcortical mechanisms.

Sleep loss has been shown to alter intracortical excitability, reducing short intracortical inhibition and intracortical facilitation in primary motor cortex (11, 12). On the other hand, spinal motoneuron excitability, as indexed by H-reflex amplitude and V-wave responses, remains largely preserved (69). Although these findings derive from total sleep deprivation protocols, they suggest that sleep pressure preferentially disrupts cortical rather than spinal motor mechanisms, making cortical dependence a plausible explanation for the dorsiflexion-specific ERD increase observed here. Viewed alongside the compensatory account, this cortical dependence offers an explanation for the movement specificity of the ERD increase: if sleep pressure reduces the efficiency of cortical processing, additional neural resource mobilization would be most evident precisely in the movement that relies most heavily on cortical drive. The absence of an analogous ERD increase during plantarflexion is therefore consistent with, and arguably predicted by, a compensatory mechanism operating at the level of cortical motor control.

### Current limitations and future directions

The present study has several limitations that bear on the interpretation of its findings. The final sample of 15 participants, after exclusions for poor EEG signal quality, is small, and effects should be regarded as preliminary pending replication in larger samples. The exclusive recruitment of young physically active males constrains generalizability: sex and age influence sleep architecture, vulnerability to sleep restriction, motor performance, and large-scale brain network organization (70–73). Whether the present findings extend to females or older adults remains an open question.

The sleep restriction protocol was implemented in participants’ home environments and monitored via actigraphy and sleep diary rather than controlled in-laboratory polysomnography. This approach has ecological validity but introduces variability in light exposure, activity, and waking behavior that cannot be fully controlled. Critically, the absence of PSG means that sleep stage composition cannot be characterized. The observed increase in sleep efficiency under CSR most likely reflects consolidated sleep under homeostatic pressure rather than improved sleep quality, and is consistent with the PVT findings, which confirm that behavioral vigilance was nonetheless impaired.

Source localization was based on a standard MNI template rather than individual structural MRIs, introducing spatial error that cannot be fully quantified. The analyses were restricted to two predefined networks, leaving open the possibility that the observed ERD increase reflects a broader cortical response to sleep pressure rather than a selective effect on sensorimotor and attentional systems. Without connectivity analyses, the parallel changes across networks remain consistent with coordinated recruitment but do not demonstrate it. Finally, the absence of a within-task attentional measure means the compensatory interpretation, while supported by the convergence of preserved performance, elevated subjective demand, and increased ERD, cannot be directly verified.

Together, these constraints mean the present findings are best regarded as hypothesis-generating. Replication in larger and more diverse samples, with PSG-verified sleep, individual head models, broader network coverage, and task-embedded attentional measures, is needed before strong mechanistic conclusions can be drawn.

## Conclusions

CSR produced a broadband increase in ERD during sustained dorsiflexion across both the sensorimotor and dorsal attention networks, without meaningful changes in motor performance. These neural changes were accompanied by impaired sustained attention and increased subjective mental demand. The movement specificity of the effect - present during the more cortically dependent dorsiflexion and absent during plantarflexion - suggests that cortically demanding movements are preferentially sensitive to sleep-related changes in cortical state. The observed pattern may reflect compensatory cortical recruitment to maintain motor output under sleep pressure, though this cannot be verified from the present data alone.

If confirmed in future work, this interpretation carries practical implications for occupational and sporting contexts in which individuals perform precise or sustained motor tasks under conditions of accumulated sleep debt. Motor output remained intact despite a higher neural cost of maintaining it, suggesting that behavioral performance alone may underestimate functional impairment.

